# The IFN response in bat cells consists of canonical and non-canonical ISGs with unique temporal expression kinetics

**DOI:** 10.1101/167999

**Authors:** Pamela C. De La Cruz-Rivera, Mohammed Kanchwala, Hanquan Liang, Ashwani Kumar, Lin-Fa Wang, Chao Xing, John W. Schoggins

**Author notes:** Corresponding author: John W. Schoggins UT Southwestern Medical Center 5323 Harry Hines Blvd. Dallas, TX 75390-9048.

## Abstract

Bats are reservoirs for a number of highly pathogenic zoonotic viruses, yet they remain relatively asymptomatic during infection. Whether this viral resistance is due to a unique innate immune system is unknown. An evolutionarily conserved feature of vertebrate antiviral immunity is the interferon (IFN) response, which triggers cellular defenses through interferon-stimulated gene (ISG) expression. While bats encode an intact IFN system, global ISG expression patterns in bat cells are not well characterized. Here, we used RNA-Seq to assess the transcriptional response to IFNα in cells derived from the bat *Pteropus alecto* (black flying fox). We show induction of more than 100 transcripts, most of which are canonical ISGs observed in other species. Kinetic gene profiling revealed that *P. alecto* ISGs fall into two unique temporal subclusters with similar early induction kinetics but distinct late-phase declines. In contrast to bat ISGs, human ISGs generally remained elevated for longer periods following IFN treatment, suggesting host-based differences in gene regulatory mechanisms. Notably, we also identified a small group of non-canonical bat ISGs, including an enzymatically active RNASEL that plays a role in controlling viral infection. These studies provide insight into the innate immune response of an important viral reservoir and lay a foundation for studies into the immunological features that may underlie unique virus-host relationship in bats.

**Significance Statement:** Bats are considered unique in their ability to resist disease caused by viruses that are often pathogenic in humans. While the nature of this viral resistance is unknown, genomic data suggest bat innate immune systems may be specialized in controlling these disease-causing viruses. A critical cell intrinsic antiviral defense system in vertebrates is the interferon response, which suppresses viral infection through induction of hundreds of interferon-stimulated genes (ISGs). In this study, we report the repertoire of ISGs and several unique features of ISG induction kinetics in bat cells. We also characterize induction and antiviral activity of bat RNASEL, which is induced by IFN in bat, but not human cells. These studies lay the foundation for discovery of potentially new antiviral mechanisms in bats, which may spur research into development of therapies to combat viral infection.

## Introduction

Bats are recognized as important viral reservoirs due to their ability to harbor a number of highly pathogenic viruses, including Nipah virus (NiV) (1), Hendra virus (HeV) (2), Marburg virus (3), SARS coronavirus (4), and Ebolavirus (5) without suffering from overt disease symptoms (6-9). A recent study identified bats as hosts for a greater proportion of zoonotic viruses than all other mammalian orders tested, with the highest viral richness found in Flavi-, Bunya- and Rhabdoviruses (10). While the mechanisms underlying disease resistance are not known, it has been speculated that bats possess unique immune components that confer innate antiviral protection (9, 11). In vertebrates, one of the first lines of defense against viral pathogens is the interferon (IFN) response. Upon viral infection, pattern-recognition receptors (PRRs) sense viral components and initiate a signaling cascade that results in the secretion of IFNs. These IFNs bind their cognate receptors to activate the JAK-STAT signaling pathway, leading to the transcriptional induction of hundreds of interferon-stimulated genes (ISGs), many of which have antiviral activity (12).

*Pteropus alecto* (also known as the black flying fox or black fruit bat) is an asymptomatic natural reservoir for the highly lethal henipaviruses (13, 14). Studies of the recently sequenced *P. alecto* genome revealed that genes for key components of antiviral immunity are conserved in bats, such as pathogen sensors (including toll-like receptors (15), RIG-I-like helicases, and NOD-like receptors), IFNs and their receptors, and ISGs (11, 16). Previous efforts to study bat-virus interactions have mainly focused on the host response to viral infection (17-20), but global transcriptional responses to type I IFN remain uncharacterized.

Since *P. alecto* harbors pathogenic human viruses and has an annotated genome, we sought to characterize the IFN-induced transcriptional response in this species. Gene expression analyses revealed that bat cells induce a pool of common ISGs. However, they also induced a small number of apparent novel ISGs, including 2-5A-dependent endoribonuclease (RNASEL). Kinetic analyses revealed that bat ISGs can be categorized into distinct groups depending on their temporal gene expression patterns. Additionally, maintenance of ISG expression over time differed between bat and human cells, suggesting distinct mechanisms of gene regulation.

## Results

### *Pteropus alecto*-derived cell lines respond to exogenous type I interferon

Previous studies have shown that exogenous IFNα and IFNγ treatment of cells from *P. alecto* can suppress *Pteropine orthoreovirus* Pulau virus (21) and Hendra virus (22), respectively. We treated immortalized *P. alecto* kidney-derived (PaKi) cells for 24h with universal IFNα, which is designed for activity across multiple species. We then infected the cells with two model GFP-expressing reporter viruses: a negative-sense single-stranded RNA rhabdovirus, vesicular stomatitis virus (VSV), and a positive-sense single stranded RNA flavivirus, yellow fever virus (YFV). We observed a dose-dependent inhibition of both viruses with IFN treatment (Fig 1A, B). VSV infection was maximally inhibited by only 50%, whereas YFV infection was suppressed completely at the highest IFN dose. In addition, we confirmed IFN-mediated dose-dependent and time-dependent induction of the canonical ISG *MX1* (Fig 1C, D) (23). These data confirm that PaKi cells are capable of mounting an antiviral response and highlight virus-specific sensitivities to IFN in this cellular background.

**Figure 1.**
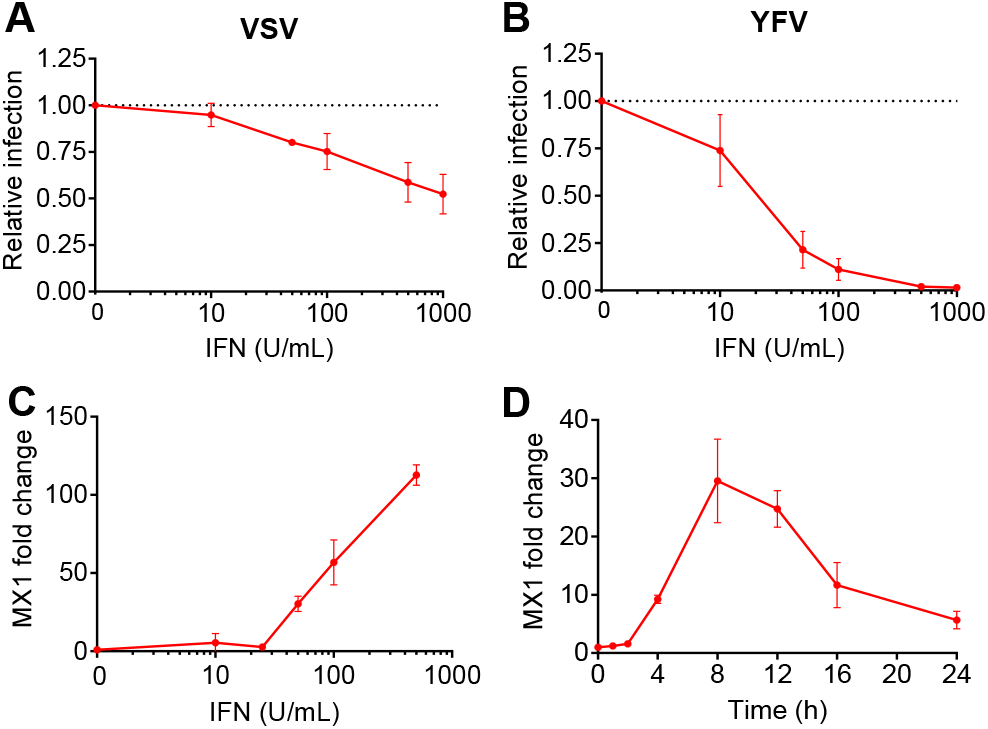
Bat-derived cells respond to Type I IFN. **(A)** PaKi cells were treated with increasing doses of IFNα for 24h and then infected with VSV-GFP at an MOI of 1. The percentage of infected cells was quantified using flow cytometry and normalized to a mock-treated control. Data are represented as mean ± SD for three independent experiments. **(B)** Same as (A), using YFV-17D Venus. **(C)** PaKi cells were treated with increasing doses of IFNα and RNA was harvested at 8h. MX1 mRNA levels were quantified using qRT-PCR and normalized to RPS11. Data are represented as mean ± SD for three independent experiments. **(D)** PaKi cells were treated with IFNα (50U/mL) and RNA was harvested at several time points. MX1 mRNA levels were measured using qRT-PCR and normalized to RPS11. Data are represented as mean ± SD for three independent experiments.

### IFN induces a classical ISG signature in PaKi cells

We next used total RNA sequencing (RNA-Seq) to profile the global transcriptional response of PaKi cells treated with IFNα over time (Fig 2A). Transcriptome assembly analysis across all experimental conditions returned approximately 30,000 genes, of which 11,559 met a minimal read count threshold (mean log_2_CPM≥0). Differential gene expression analysis revealed that IFN induced 93 genes at 4h, 104 at 8h, 103 at 12h, and 103 at 24h (fold change (FC)≥1.5, FDR≤0.05) (Fig 2B). There were no downregulated genes at 4h, 2 at 8h, 105 at 12h, and 279 at 24h. However, statistical significance of upregulated genes was more robust than the statistical significance of the downregulated genes. Overall, IFN treatment of PaKi cells produced a positive gene induction signature.

**Figure 2.**
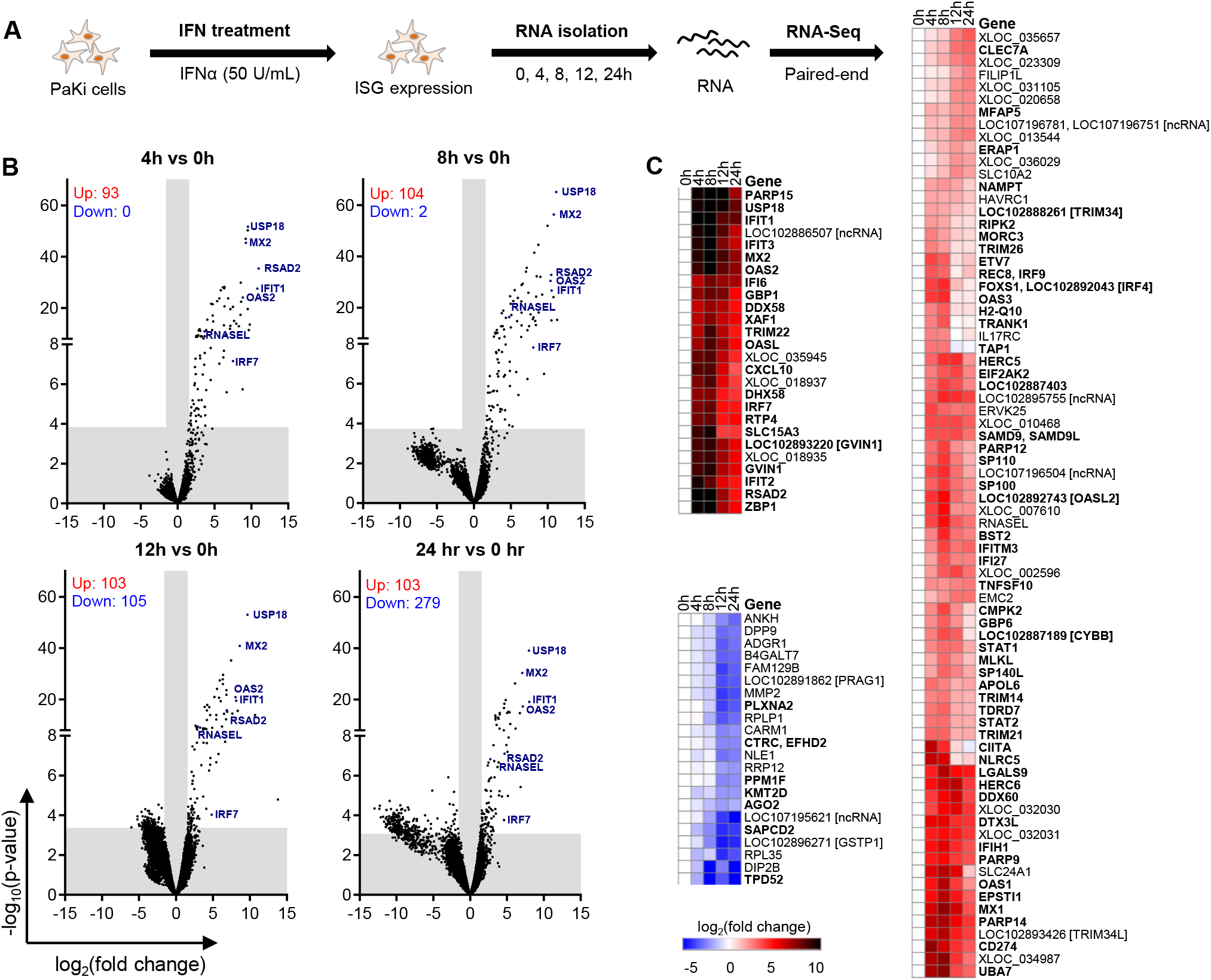
PaKi transcriptome response to IFN. **(A)** Experimental schematic. **(B)** Volcano plots for all time points. Differentially-expressed genes are quantified on the top left. Grey bars indicate log_2_FC≥1.5 (vertical) or FDR≤0.05 (horizontal). Data is a result of three independent experiments **(C)** Heatmap including genes that meet the cutoffs mean log_2_CPM≥0, log_2_FC≥2, FDR≤0.05 for at least 2 time points. XLOC genes represent unannotated genes.

Next, heat maps were generated to assess individual gene induction, using a cutoff of FDR≤0.05 and FC≥4 for at least 2 time points (Fig 2C). Many genes in this list have known roles in innate immunity, including well-known ISGs *(IFIT1, MX1, OAS2, RSAD2*/viperin, *USP18*), members of the JAK-STAT signaling cascade (*STAT1, STAT2, IRF9*), pattern recognition receptors (*DDX58*/RIG-I, *IFIH1*/MDA5, *ZBP1*) and transcription factors (*ETV7, IRF7, SP110*) (24). Notably, we detected induction of *GVIN1* (interferon-induced very large GTPase), which is predicted to encode a protein in bats but is annotated as a pseudogene in humans. This list also included transcripts predicted to encode an endogenous retrovirus *(ERVK25)* and several transcripts predicted to be long non-coding RNAs.

Differentially-expressed genes were cross-referenced to the INTERFEROME v2.0 database (25) to determine if they had previously been reported as IFN-induced genes. At the 4h and 8h time points, more than 80% of the genes in our data set overlapped with INTERFEROME v2.0. Since the INTEFEROME database consists predominantly of human and mouse data sets, this result suggests that antiviral responses in bat cells include a conserved repertoire of IFN-inducible genes commonly found in other mammalian species.

### Differential temporal regulation of *P. alecto* ISGs

We next used a clustering algorithm to group genes in the RNA-Seq data set based on induction kinetics, without imposing fold change or FDR cutoffs (26). This analysis revealed that genes were organized into eight subclusters (SC) based on changes in expression levels throughout the IFN time course (Fig 3A). Genes in SC1 and SC2 increased in expression after 8h. Genes in SC3 and SC4 were induced by 4h and peaked at 4-8h, with genes in SC3 returning close to baseline by 12h and SC4 genes remaining elevated for at least 24h. These two SCs were highly-enriched for genes found in the INTERFEROME v2.0 database. Genes in SC5 increased slightly and peaked at 8h, returning to baseline levels by 12h. Levels of SC6 genes either increased or decreased by 4h, followed by decreased levels at 12-24h. SC7 gene expression levels decreased sharply by 8h, followed by a partial recovery by 12h and further decrease in expression by 24h. Finally, genes in SC8 exhibited a sustained decrease in expression levels starting at 8-12h. These data demonstrate that interferon induces distinct subsets of genes which are characterized by differing temporal expression patterns.

**Figure 3.**
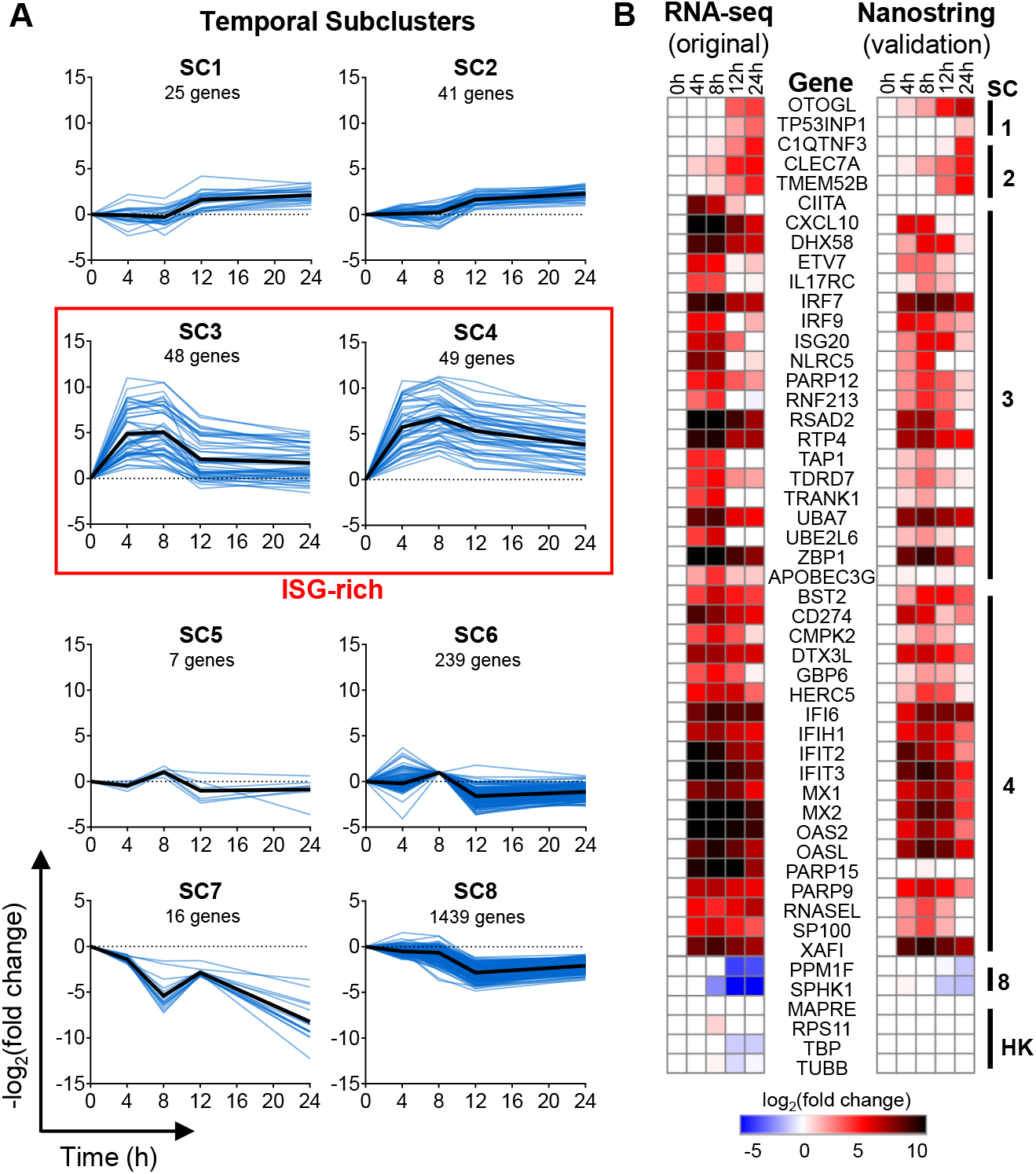
Temporal clustering analysis. **(A)** Temporal subclusters. The number of genes in each subcluster is indicated below the title. Blue lines represent mean log_2_(fold change) over time for a single gene. Black lines represent median of all genes in the subcluster. **(B)** RNA-seq validation using Nanostring. RNA was isolated from PaKi cells and gene expression over time was compared between RNA-seq data (left) and Nanostring data (right). Data presented is the mean of three independent experiments. HK = housekeeping genes.

### Orthogonal validation of RNA-Seq data

To validate our RNA-seq results, we used Nanostring nCounter technology, which uses colorimetrically-barcoded DNA probes for direct detection of mRNA without the use of nucleic acid amplification. A customized nCounter codeset was designed containing 50 genes from several temporal SCs, with a focus on the ISG-rich SC3 and SC4 (Fig S2). Temporal expression profiles using Nanostring were generally similar to the RNA-Seq data (Fig 3B). SC1 and SC2 contained genes with increased expression levels at the later time points. SC3 and SC4 had genes with peak expression levels at 8h, with genes in SC4 exhibiting expression levels that remained elevated at 24h. Genes in SC8 decreased in expression over time, with the lowest expression levels observed at 24h.

### Human vs Bat Temporal ISG Regulation

We next used Nanostring to compare temporal regulation of ISGs between bat and human cell lines. We first compared gene expression between PaKi and HEK293 as both cell lines are kidney-derived, but HEK293 cells responded poorly to type I IFN (Fig S1). After screening for robust IFN responses in other human cell lines, we chose human A549 cells for comparative studies (Fig 4A). A striking difference was observed between expression profiles of SC1 and SC2 when comparing PaKi and A549 cells. Of the 5 selected genes in SC1 and SC2, none were significantly upregulated in IFN-treated A549 cells. Similarly, the decreased expression levels observed for bat genes in SC8 were not observed in A549 cells.

**Figure 4.**
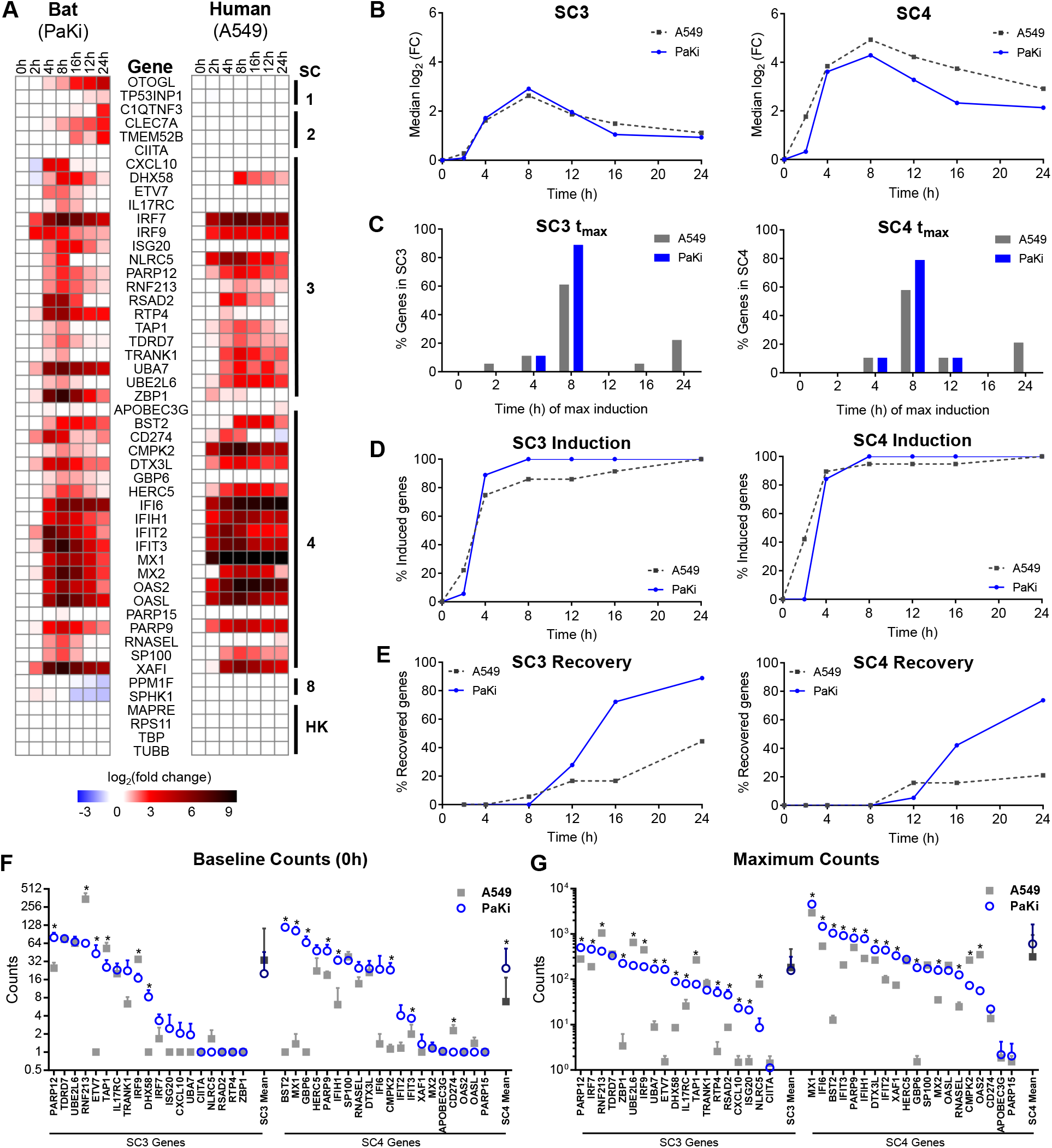
ISG expression over time in bat and human cells. **(A)** Nanostring gene expression for PaKi (left) and A549 cells (right). PaKi data is the same shown in Fig 3B, with additional time points. Data presented is the mean of three independent experiments. HK = housekeeping genes. **(B)** Expression over time for PaKi and A549 genes in SC3 and SC4. Each line represents the median log_2_(FC) for all genes in a SC. **(C)** Time at peak induction for PaKi and A549 genes in SC3 and SC4. **(D)** Percent genes induced to levels ≥50% of maximum induction at specified time points in SC3 and SC4. **(E)** Percent genes that recovered to levels ≤50% of maximum induction at specified time points in SC3 and SC4. **(F)** Baseline mRNA counts for PaKi and A549 genes in SC3 and SC4. Data are represented as mean + SD for three independent experiments. Darker symbols represent mean counts for each subcluster. * = p-value <0.05 using Student’s t-test. **(G)** Maximum mRNA counts for PaKi and A549 genes in SC3 and SC4. Data are represented as mean + SD for three independent experiments. Darker symbols represent mean counts for each subcluster. * = p-value <0.05 using Student’s t-test.

For further analysis, we chose to focus on previously-reported ISGs, particularly those found in SC3 and SC4. When comparing changes in gene expression in SC3, the median fold-change of all genes in SC3 followed similar trends between bat and human cell lines (Fig 4B). In SC4, however, genes from A549 cells exhibited higher fold induction and remained elevated at later time points when compared to PaKi cells.

We next determined the time point at which we observed maximum gene induction (t_max_). For both SC3 and SC4, approximately 80% of IFN-induced PaKi genes peaked at 8h (Fig 4C). However, A549 genes had a bimodal pattern within both SCs, with most genes peaking at 4-8h and a smaller subset peaking at 12h or later.

To identify potential differences in induction kinetics, we calculated the percentage of genes that were induced to at least 50% of maximum expression for each time point. Greater than 80% of PaKi genes in both SCs were induced by 4h (Fig 4D). A549 genes in SC3 behaved similarly, although a small subset of genes was induced by 2h. However, approximately 40% of A549 genes in SC4 were induced by 2h, indicating faster induction of this subset of genes. Genes in this list include *HERC5, IFI6, IFIH1, MX1, NLRC5, OAS2, OASL*, and *PARP12*. Notably, PaKi cells express higher baseline levels of ISGs such that by full induction at 8h, PaKi cells express similar or greater mRNA counts compared to A549 cells, despite faster induction in A549 cells (Fig 4D, Fig 4G, and Fig S1).

Next, we calculated the percentage of genes that had recovered to below 50% of maximum expression for each time point. In both SCs, recovery occurred earlier in PaKi cells, with most genes recovering by 16h (Fig 4E). In contrast, the expression levels for most A549 genes remained elevated throughout the time course. Together, these data suggest that the regulatory mechanisms governing IFN-mediated gene induction and maintenance of gene expression differs in each cell type, particularly with genes in SC4.

It was recently reported that *P. alecto* tissues have constitutive and ubiquitous expression of IFNα, suggesting that bat cells may be primed to inhibit viral infection due to constitutive expression of certain ISGs (21). Indeed, we observed overall higher ISG mRNA levels in unstimulated PaKi cells, particularly the SC4 genes (Fig 4F). In addition, we found that over half of SC4 PaKi ISGs were induced to higher maximum counts than corresponding A549 ISGs (Fig 4G).

Bat cells express multiple non-canonical ISGs, including an active RNASEL. Our gene expression profiling revealed several genes not previously reported to be ISGs. These included *EMC2, FILIP1, IL17RC, OTOGL, SLC10A2*, and *SLC24A1* (Fig 2A). In addition, we observed IFN-mediated induction of *RNASEL*, which encodes a 2’-5’-oligoadenylate synthetase-dependent ribonuclease. This protein exerts its antiviral effect by degrading viral RNA in response to 2’-5’-linked oligoadenylates, which are generated by the IFN-inducible oligoadenylate synthase (OAS) family of enzymes upon stimulation by dsRNA in the cytosol (27). In humans, RNASEL is a constitutively-expressed latent enzyme and is not IFN-inducible (Fig 4A, Fig S1) (28). Interestingly, we observed a dose-dependent induction of *RNASEL* in IFN-treated PaKi cells (Fig 5A). Of note, similar mRNA expression levels of *RNASEL* were observed in unstimulated human and bat cell lines (Fig 4F and Fig S1). In addition, we observed IFN-mediated *RNASEL* induction in brain-derived (PaBr) and lung-derived (PaLu) *P. alecto* cell lines (Fig 5B), suggesting IFN-mediated induction of RNASEL is not cell type-specific.

**Figure 5.**
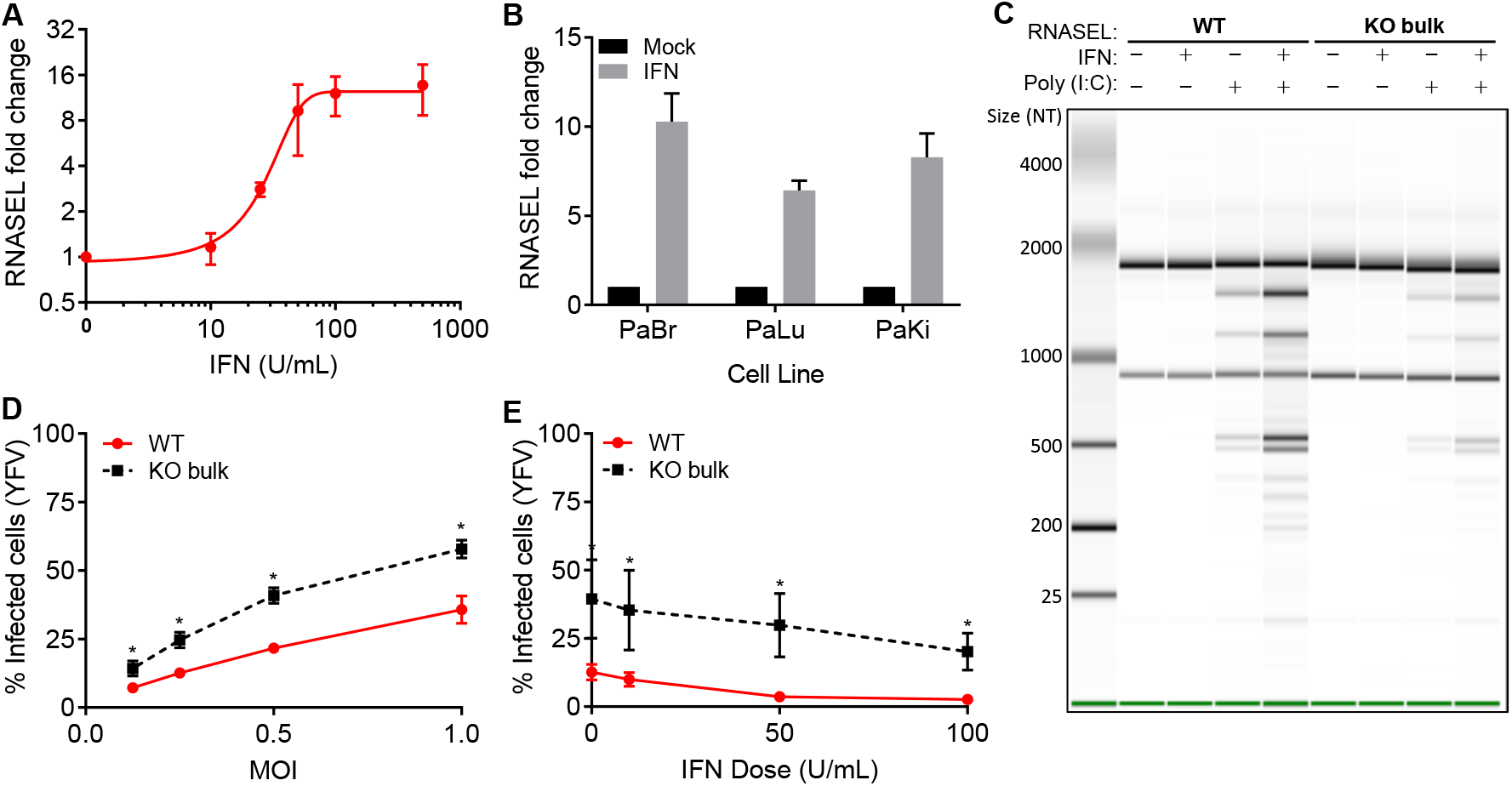
RNASEL is IFN-induced in *P. alecto*. **(A)** PaKi cells were treated with increasing doses of IFNα and RNA was harvested at 8h. RNASEL mRNA levels were measured using qRT-PCR and normalized to untreated control using RPS11 as a housekeeping control. Data are represented as mean ± SD for two independent experiments. **(B)** RNASEL induction in *P. alecto* brain, lung, and kidney cells. Cells were treated with IFNα (100U/mL) and RNA was harvested at 8h. RNASEL levels were quantified using qRT-PCR and normalized to untreated control using RPS11 as housekeeping control. Data are represented as mean ± SD for three independent experiments. **(C)** PaKi cells were treated with IFNα (100U/mL) or mock-treated for 24h, followed by transfection with poly(I:C) (100ng/mL) or mock-transfected for 4h and RNA integrity was monitored using a bionalyzer. 100 ng total RNA was run per lane. Data is representative of two independent experiments. **(D)** WT or RNASEL KO bulk PaKi cells were infected with YFV-17D Venus for 24h and viral infection was quantified using flow cytometry. Data are represented as mean ± SD for three independent experiments. * = p-value < 0.05 using Student’s t-test. **(E)** WT or RNASEL KO bulk PaKi cells were treated with increasing doses of IFNα for 24h, then infected with YFV-17D Venus at an MOI of 0.5 for 24h. Viral infection was quantified using flow cytometry. Data are represented as mean ± SD for three independent experiments. * = p-value < 0.05 using Student’s t-test.

Next, we constructed RNASEL-deficient PaKi cells using CRISPR (29). Due to lack of a bat-specific RNASEL antibody, we were not able to monitor RNASEL protein levels. However, we detected modifications to the *RNASEL* gene in the PaKi knock-out bulk population (KO bulk) as compared to the parental wild-type (WT) population (Table S1). To test if RNASEL protein was functional, we activated RNASEL with poly (I:C) transfection after IFN or mock treatment and monitored RNA integrity (Fig 5C) (30, 31). We observed rRNA degradation when cells were treated with poly(I:C), but not with IFN alone. Treating cells with IFN for 24h to induce RNASEL expression before poly (I:C) transfection resulted in increased RNA degradation and accumulation of smaller products, suggesting increased RNASEL activity. The RNA degradation observed in the WT cells was reduced in the bulk KO cells. Together, these data indicate that bat RNASEL is a functional ribonuclease that, unlike its human ortholog, is IFN-inducible.

To test if the presence of RNASEL is important in the context of viral infection, we infected both WT and bulk KO cells with YFV17D-Venus and quantified infectivity after one viral life cycle (Fig 5D). RNASEL KO cells were more permissive to infection at all doses used, suggesting that RNASEL is important for suppression of initial viral infection. To test if RNASEL induction played a role in the protective effect of the IFN response, we treated PaKi WT and KO bulk cells with increasing doses of IFNα for 24h, then infected with YFV17D-Venus (Fig 5E). Consistent with previous results, KO bulk cells were more permissive to infection than WT cells. In addition, KO bulk cells were resistant to the protective activity of IFN. IFNα (100U/mL) pre-treatment resulted in 80% reduction of infection in WT cells, but only 50% reduction in bulk KO cells. Together, these data suggest RNASEL contributes to inhibition of viral infection in bat cells, particularly in the context of the IFN response.

## Discussion

This study aimed to identify IFN-stimulated transcripts in the viral reservoir *P. alecto*. Transcriptional analysis revealed over 100 genes induced in response to IFNα. Most of these genes have been previously described as ISGs, suggesting strong evolutionary conservation of the ISG pool, as would be predicted by previous genomic studies of immune genes in *P. alecto* (11, 16).

We have provided a framework of *P. alecto* ISG induction, organized by early, mid, and late responses to type I IFN. Our data set reveal a previously unappreciated subtlety in the temporal induction and regulation of ISG expression. A recent study of IFN-inducible gene kinetics in mouse cells reported a profile that resembles a combination of our SC3 and SC4 data sets (32). Our study further refines this kinetic analysis by delineating two separate ISG pools based on unique temporal induction profiles. While both SCs are characterized by similar peak mRNA levels and a subsequent decline by 12-24 hours, SC4 contained some genes that remained elevated. These genes may offer residual antiviral protection, even when IFN signaling has returned to basal levels. In addition, many genes in SC4 have both higher baseline and higher maximal induction levels in bat cells compared to human cells, which could contribute to species-specific differences in susceptibility to viral infection. Additional studies are needed to explore the mechanisms underlying unique expression of certain ISGs in bats cells. Compared to human cells, bat cells have a more rapid decline in ISG levels, suggesting tightly-regulated expression kinetics. The reason for this strict transcriptional regulation remains unclear, but such a mechanism may exist to prevent excessive inflammation in a highly metabolically active host (11).

We also identified several previously unrecognized, or non-canonical ISGs, including RNASEL. The OAS/RNASEL pathway consists of OAS proteins that recognize viral dsRNA and subsequently catalyze the formation of short oligoadenylates that act as second messengers to activate the constitutively-expressed latent enzyme RNASEL, which degrades viral genetic material. Due to cleavage of both viral and cellular RNA, RNASEL activation can also lead to apoptosis of infected cells. In addition, the short RNA fragments created by RNASEL can potentiate the IFN response by activating the cytosolic RNA sensor RIG-I (33). The induction of RNASEL in response to IFN in bats may provide an additional layer of antiviral protection. Indeed, knockout of RNASEL increased viral susceptibility of *P. alecto-derived* cells. Although RNASEL induction itself does not result in significant nuclease activity, stimulation with poly (I:C) is sufficient to cause degradation of total RNA in the cell. Unlike in humans, where only the upstream OAS proteins are IFN-induced, bat cells co-induce both components of the OAS/RNASEL pathway, likely creating a more rapid and potent effect that would inhibit viral replication before extensive viral spread could occur. RNASEL induction may also be a counterdefense against viruses that subvert the pathway through through various immune evasion mechanisms, such as degradation of RNASEL by L* protein of murine Theiler’s virus(34), or increased expression of an RNASEL inhibitor by HIV(35) and EMCV(36).

Additional studies will be needed to determine whether these non-canonical bat ISGs contribute to unique viral resistance in bats. It is also possible that canonical bat ISGs have evolved unique effector functions that contribute to reduced viral susceptibility. Uncovering mechanisms of bat ISGs will provide insight into the innate immune responses of an important viral reservoir and may inform research and development of antiviral therapies.

## Methods & Materials

### Cell lines

*P. alecto*-derived PaBr (brain), PaLu (lung), and PaKi (kidney) immortalized cell lines (37) were grown at 37°C and 5% CO_2_ and passaged in DMEM/F-12 media (GIBCO) supplemented with 10% FBS. Human A549 lung adenocarcinoma and HEK293 cells were grown at 37°C and 5% CO_2_ and passaged in DMEM media (GIBCO) supplemented with 10% FBS and 1X non-essential amino acids (NEAA) (GIBCO #11140076).

### Viruses

YFV-Venus (derived from YF17D-5C25Venus2AUbi) stocks were generated by electroporation of in vitro-transcribed RNA into *STAT1*^-/-^ fibroblasts as previously described (12). VSV-GFP (provided by Jack Rose) was generated by passage in BHK-J cells. For both viruses, virus-containing supernatant was centrifuged to remove cellular debris and stored at -80C until use.

### Viral infection

Cells were seeded into 24-well plates at a density of 1x10^5^ cells per well. Viral stocks were diluted into DMEM/F-12 media supplemented with 1% FBS to make infection media. Media was aspirated and replaced with 200 μL of infection media. Infections were carried out at 37°C for 1h, then 800 μL DMEM/F-12 media supplemented with 10% FBS was added back to each well. After approximately one viral life cycle, cells were harvested for flow cytometry.

### Flow cytometry

Cells were detached using Accumax, fixed in 1% PFA for 10 min at room temperature, and pelleted by centrifugation at 800xg. Fixed cell pellets were resuspended in 200 μL FACS buffer (1X PBS supplemented with 3% FBS). Samples were run on a Stratedigm S1000 instrument using CellCapTure software and gated based on GFP fluorescence. Data analysis was done using FlowJo software (v9.7.6).

### Interferon treatment and RNA isolation

Cells were seeded into 6-well plates at 3x10^5^ cells per well. The following day, cells were treated with 2 mL of DMEM F-12 media supplemented with 10% FBS and 50 units/mL of Universal Type I IFN Alpha (PBL Assay Science Cat. #11200). The cells were harvested by aspirating the media, washing twice with 2 mL of sterile 1X PBS, then lysing with 350 μL RLT buffer from the RNeasy Mini kit (Qiagen). The cell lysate was stored at -80C until RNA isolation was completed using the RNeasy kit following the manufacturer’s protocol.

### RNA-Seq

Total RNA samples for each time point were prepared in three independent experiments as described above. The Agilent 2100 Bioanalyzer was used to determine RNA quality; only samples with a RIN Score >9 were used. A Qubit fluorometer was used to determine RNA concentration. One μg of DNAse-treated total RNA was prepared with the TruSeq Stranded Total RNA LT Sample Prep Kit (Illumina). rRNA-depleted total RNA was fragmented and used for strand-specific cDNA synthesis. cDNA was then A-tailed and ligated with indexed adapters. Samples were then PCR amplified, purified with AmpureXP beads, and re-validated on the Agilent 2100 Bioanalyzer and Qubit.

Normalized and pooled samples were run on the Illumina HiSeq 2500 using SBS v3 reagents. Paired-end 100 bp read length Fastq files were checked for quality using fastqc (http://www.bioinformatics.babraham.ac.uk/projects/fastqc) and fastq_screen (http://www.bioinformatics.babraham.ac.uk/projects/fastqscreen/) and were quality trimmed using fastq-mcf (https://github.com/ExpressionAnalysis/ea-utils/blob/wiki/FastqMcf.md). Trimmed fastq files were mapped to Pteropus alecto (black flying fox) assembly ASM32557v1 (ftp://ftp.ncbi.nih.gov/genomes/Pteropusalecto) using TopHat (38). Duplicates were marked using picard-tools (https://broadinstitute.github.io/picard/). Reference annotation based transcript assembly was done using cufflinks (39), and read counts were generated using featureCounts (40). Pairwise differential expression analysis was performed using edgeR (41), and longitudinal analysis was performed using time course (26) after data transformation by voom (42). Differentially-expressed unannotated genes were manually annotated using a nucleotide BLAST (43) search for homologous genes (indicated with an asterisk).

### Heatmaps

Heatmaps were constructed using GENE-E and Morpheus, both available at https://software.broadinstitute.org.

### Nanostring analysis

Total RNA was isolated from IFN-treated samples as described above. 100 ng of total RNA in 5 μL was used as input in each reaction for NanoString assay. A customized panel containing both *P. alecto* and *H. sapiens* genes (Schoggins_1_C4066, NanoString Technologies, Seattle, WA, USA) was used to measure the expression of 100 genes in the RNA samples by following the NanoString nCounter XT Codeset Gene Expression Assay manufacturer’s protocol. Following hybridization, transcripts were quantitated using the nCounter Digital Analyzer.

### qRT-PCR

Total RNA was prepared as described above. Reactions were prepared with the QuantiFast SYBR Green RT-PCR kit (Qiagen #204154), using 50 ng total RNA per reaction. Samples were run on the Applied Biosystems 7500 Fast Real-Time PCR System using 7500 Software v2.0.6. Primers used to amplify RNASEL are 5’-CCACCCTGGGGAAAATGTGA-3’ and 5’-GGAGGATCCTGCTTGCTTGT-3’. Primers used to amplify housekeeping gene RPS11 are 5’-ATCCGCCGAGACTATCTCCA-3’ and 5’-GGACATCTCTGAAGCAGGGT-3’.

### DNA constructs and plasmid propagation

lentiCRISPR v2 was a gift from Feng Zhang (Addgene plasmid # 52961). RNASEL targeting guides were generated by cloning annealed, complementary 20-bp oligos with *Esp3I*-compatible overhangs (5’-caccgAGACCCACACCCTCCAGCAG-3’ and 5’-aaacCTGCTGGAGGGTGTGGGTCTc-3’) targeting the *P. alecto RNASEL* gene into the lentiCRISPRv2 backbone as described in (44). CRISPR guide oligos were designed using CRISPRdirect (45). Proper assembly was confirmed using Sanger sequencing.

### Lentiviral pseudoparticles

Lentiviral pseudoparticles were made as described in (46), with some modifications. Briefly, 2x10^6^ HEK293T cells were seeded on a poly-lysine coated 10 cm plate. The following day, media was changed to 7.5 mL DMEM with 3%FBS and 1x NEAA. Cells were co-transfected with 5 μg lentiCRIPSRv2, 2.5 μg pCMV-VSVg and 3.5 μg pGag-pol. 4h post-transfection, media was changed to 7.5 mL fresh DMEM with 3% FBS and 1x NEAA. Supernatant was collected 48h post-transfection, filtered through 0.45 μm to remove debris, and stored at -80°C until use.

### *RNASEL* KO bulk PaKi cell lines

3x10^5^ PaKi cells were seeded on 6-well plates. The following day, media was changed to DMEM/F-12 supplemented with 3% FBS, 4 μg/mL polybrene and 20 mM HEPES. LentiCRISPRv2 lentiviral pseudoparticles were added and cells were spinoculated at 800xg for 45 min at 37°C. Media was changed to DMEM/F-12 with 10% FBS immediately following spinoculation. 48h after transduction, cells were pooled into a 10-cm dish and selected in DMEM/F-12 with 10% FBS and 5 μg/mL puromycin.

### Genomic characterization of PaKi *RNASEL* KO bulk cell lines

Genomic DNA was isolated from wild-type or RNASEL KO bulk populations using the DNeasy Blood and Tissue Kit (Qiagen). The region falnking the CRISPR-targeted sequence was amplified via PCR using primers (5′-ATGGAGACCAAGAGCCACAACA-3′) and (5’-CGTCCTCGTCCTGGAAATTGA-3’). PCR products were gel-purified using the QIAquick Gel Extraction Kit (Qiagen) and subsequently used in TOPO cloning reactions using the TOPO TA Cloning Kit (Thermo Fisher). Several colonies were selected for each cell background and colony PCR was used to amplify the CRISPR-targeted region. Samples were then analyzed using Sanger sequencing.

### rRNA degradation assay

3x10^5^ PaKi cells were seeded on 6-well plates. The following day, universal IFN (100 U/mL) was added to the media. After 24h, cells were transfected with 100 ng/mL poly(I:C) in Optimem using Lipofectamine 3000. After 4h, RNA was harvested using the RNeasy Mini Kit (Qiagen) and RNA integrity was measured on a Total RNA Nano chip using an Agilent 2100 Bioanalyzer.

## Acknowledgements

We would like to thank Jeanine Baisch and Cynthia Silverman from the Genomics Core at Baylor Research Institute for assistance with Nanostring samples, and the McDermott Center Sequencing Core and Bioinformatics Core for sequencing and analysis. This work was supported in part by NIH grant AI117922 (J.W.S.), the UT Southwestern Endowed Scholars program (J.W.S.), the UT Southwestern High Impact / High Risk Grant Program (J.W.S.), the William F. and Grace H. Kirkpatrick Award (P.C.D.) and the NRF-CRP grant NRF2012NRF-CRP001-056 (L-F.W). C.X. was partially supported by NIH grant UL1TR001105.

---

**Figure S1.**
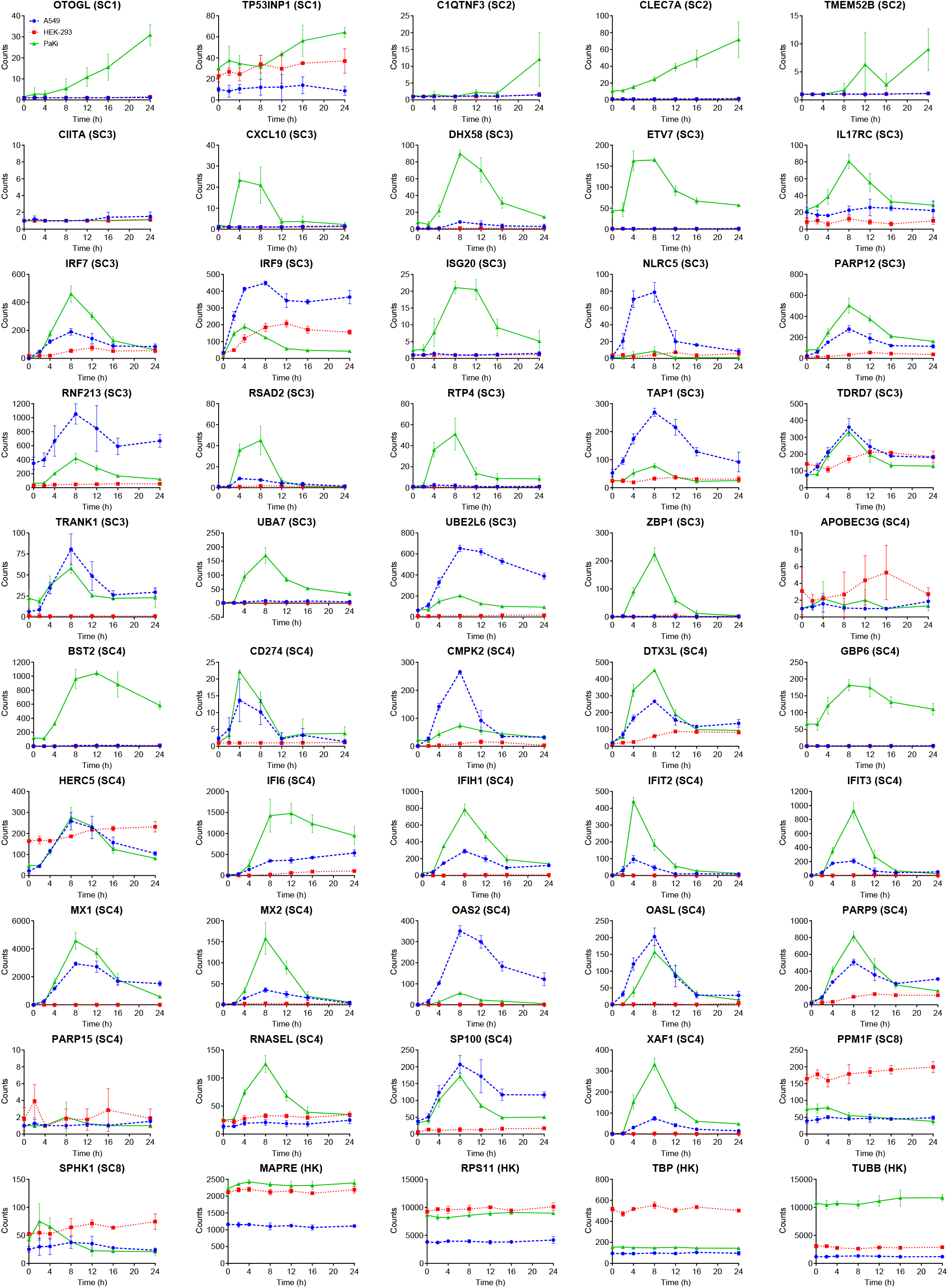
Normalized Nanostring counts over time in bat and human cells. A549 (blue), HEK293 (red), or Paki (green) cells were treated with IFNα (50U/mL) and RNA was harvested at the indicated time points. A customized Nanostring nCounter library was used to detect mRNA counts of 50 genes. Data are represented as mean ± SD for three independent experiments.

